# SEM: sized-based expectation maximization for characterizing nucleosome positions and subtypes

**DOI:** 10.1101/2023.10.17.562727

**Authors:** Jianyu Yang, Kuangyu Yen, Shaun Mahony

**Affiliations:** Center for Eukaryotic Gene Regulation, Department of Biochemistry and Molecular Biology, Pennsylvania State University, University Park, PA, USA; State Key Laboratory of Experimental Hematology, National Clinical Research Center for Blood Diseases, Haihe Laboratory of Cell Ecosystem, Institute of Hematology & Blood Diseases Hospital, Chinese Academy of Medical Sciences & Peking Union Medical College, Tianjin 300020, China; Department of Developmental Biology, School of Basic Medical Sciences, Southern Medical University, Guangzhou, China

## Abstract

Genome-wide nucleosome profiles are predominantly characterized using MNase-seq, which involves extensive MNase digestion and size selection to enrich for mono-nucleosome-sized fragments. Most available MNase-seq analysis packages assume that nucleosomes uniformly protect 147bp DNA fragments. However, some nucleosomes with atypical histone or chemical compositions protect shorter lengths of DNA. The rigid assumptions imposed by current nucleosome analysis packages ignore variation in nucleosome lengths, potentially blinding investigators to regulatory roles played by atypical nucleosomes.

To enable the characterization of different nucleosome types from MNase-seq data, we introduce the Size-based Expectation Maximization (SEM) nucleosome calling package. SEM employs a hierarchical Gaussian mixture model to estimate the positions and subtype identity of nucleosomes from MNase-seq fragments. Nucleosome subtypes are automatically identified based on the distribution of protected DNA fragment lengths at nucleosome positions. Benchmark analysis indicates that SEM is on par with existing packages in terms of standard nucleosome-calling accuracy metrics, while uniquely providing the ability to characterize nucleosome subtype identities.

Using SEM on a low-dose MNase H2B MNase-ChIP-seq dataset from mouse embryonic stem cells, we identified three nucleosome types: short-fragment nucleosomes, canonical nucleosomes, and dinucleosomes. The short-fragment nucleosomes can be divided further into two subtypes based on their chromatin accessibility. Interestingly, the subset of short-fragment nucleosomes in accessible regions exhibit high MNase sensitivity and display distribution patterns around transcription start sites (TSSs) and CTCF peaks, similar to the previously reported “fragile nucleosomes”. These SEM-defined accessible short-fragment nucleosomes are found not just in promoters, but also in enhancers and other regulatory regions. Additional investigations reveal their co-localization with the chromatin remodelers Chd6, Chd8, and Ep400.

In summary, SEM provides an effective platform for distinguishing various nucleosome subtypes, paving the way for future exploration of non-standard nucleosomes.

## INTRODUCTION

The nucleosome is the basic packaging unit of chromatin, typically comprising ∼147bp DNA wrapped around the histone octamer ^1^. Nucleosomes participate in gene regulation by both physically impeding access to DNA and by serving as a substrate for interactions with regulatory proteins ^2^. Chemical modifications on histone tails, including methylation, acetylation, and ubiquitination, can alter DNA affinity to the octamer and can be engaged by regulatory proteins ^3,4^. Several histone tail chemical modifications are well correlated with transcriptional activity or repression ^5–7^. Histone variants, such as H2A.Z and H3.3, have also been reported to play roles in many important biological events, such as DNA replication and enhancer activity ^8,9^.

The most common technique for studying the nucleosome landscape across the genome is micrococcal nuclease sequencing (MNase-seq). MNase is an endo-exonuclease that preferentially digests accessible DNA between nucleosomes. After size selection, mono-nucleosome-sized DNA fragments are retained for high-throughput sequencing ^10,11^. However, depending on the composition of the nucleosome and the factors engaging the nucleosome, not all nucleosomes protect the canonical 147bp of DNA. For example, nucleosomes engaged by Pol II lose one H2A-H2B dimer and transiently become hexamers ^12^. Nucleosomes containing the histone variant H2A.Z can exhibit a distinct unwrapping state from the canonical nucleosome ^13^. Some studies employing alternative nucleosome mapping methods, including low-dose MNase-seq, methidiumpropyl-EDTA sequencing (MPE-seq) ^14^, Cleavage Under Targets and Release Using Nuclease (CUT&RUN) ^15^, and chemical mapping ^16^, found MNase-sensitive nucleosome subtypes in yeast and mouse NFRs which protect shorter DNA fragments than canonical nucleosomes ^14–16^. These studies have begun revealing variations in nucleosome composition across the genome.

Despite experimental results that have characterized a wider diversity of nucleosome composition, most nucleosome-calling software packages still assume that nucleosomes uniformly protect ∼147bp of DNA ^17–19^. Current approaches use this rigid assumption when estimating the locations and occupancy properties of nucleosomes, making their performance sub-optimal when characterizing nucleosomes of non-canonical DNA length. Although some packages have begun incorporating the ability to detect nucleosome position dynamics ^17,18^, none are yet able to distinguish various nucleosome subtypes from MNase-seq data.

To resolve the lack of an effective method for characterizing nucleosome subtypes, we introduce a new nucleosome-calling package called Size-based Expectation Maximization (SEM). SEM first analyzes the overall fragment size distribution to determine which types of nucleosomes are detectable within a given MNase-seq dataset. SEM then employs a hierarchical Gaussian mixture model to accurately estimate the locations and occupancy properties of nucleosomes and to assign subtype identities to each detected nucleosome. Thus, in addition to reporting the properties often considered by common nucleosome-calling packages, including nucleosome dyad location, occupancy, and fuzziness, SEM also distinguishes various nucleosome subtypes from a single MNase-seq experiment.

To demonstrate SEM’s ability to distinguish nucleosome subtypes, we apply it to a low dose MNase-ChIP-H2B dataset from mouse embryonic stem cells (mESCs) ^14^. The MNase-ChIP-seq dataset retained the full fragment size range by omitting mono-nucleosome size selection. On this low-dose MNase-ChIP-H2B data, SEM identifies three nucleosome types, including short-fragment nucleosomes, canonical nucleosomes, and di-nucleosomes. Downstream analysis shows that short-fragment nucleosomes located within accessible regions share similar properties to the so-called “fragile” nucleosomes reported in previous publications ^14,16^. Investigation of these SEM-defined accessible short-fragment nucleosomes genome-wide shows that they are not exclusively restricted to transcription start sites (TSSs) and CTCF binding sites. We further find associations between accessible short-fragment nucleosomes and specific chromatin remodelers. Exploring the remaining short-fragment nucleosomes that lie outside of accessible regions suggests that they are the result of MNase digestion bias on AT-rich DNA at nucleosome entry and exit sites. Overall, our findings demonstrate the ability of SEM to characterize various nucleosome subtypes genome-wide.

## RESULTS

### A hierarchical Gaussian Mixture Model for characterizing nucleosome types

SEM is a hierarchical Gaussian Mixture Model (GMM), which probabilistically models the positions, occupancy, fuzziness, and subtype identities of nucleosomes from MNase-seq data (**Fig. 1**). The components of the mixture model represent individual nucleosomes; the properties of each nucleosome are modeled based on the mapped locations and lengths of MNase-seq fragments. Specifically, each nucleosome component is defined by its dyad location, occupancy, fuzziness, and the probability of belonging to each nucleosome subtype.

**Figure 1:**
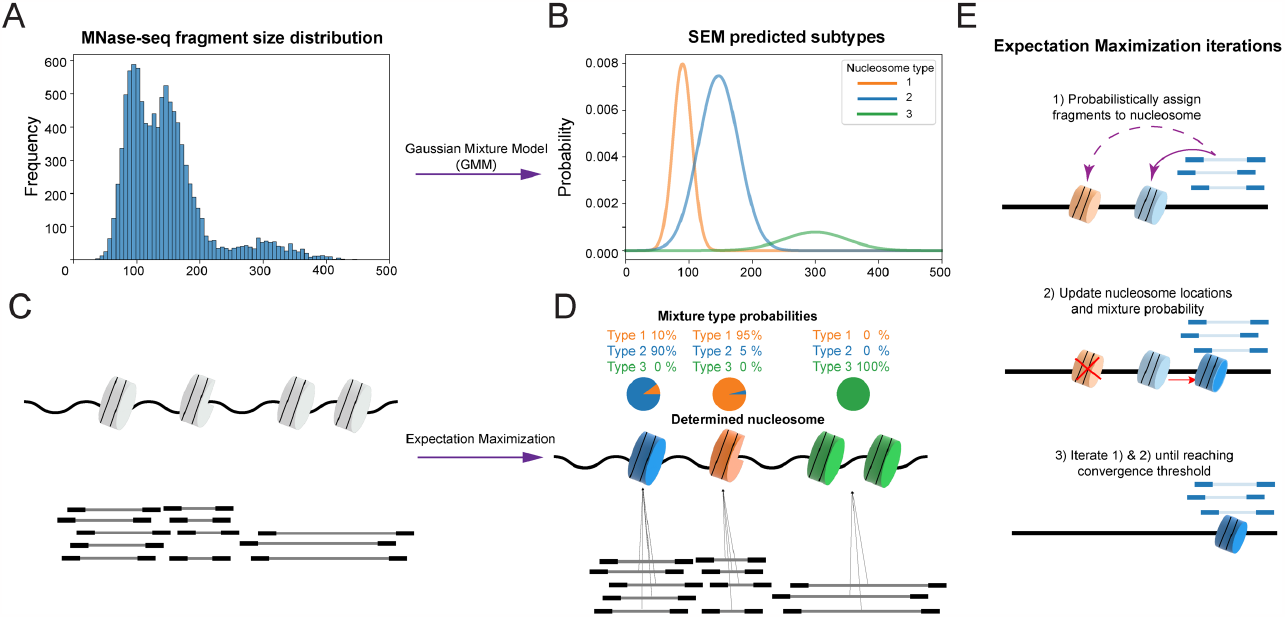
Overview of the SEM algorithm. **A)** SEM first calculates the global fragment size distribution from the MNase-seq data. **B)** A Gaussian Mixture Model is used to deconvolve the fragment size distribution into a set of nucleosome subtypes. **C)** In the Expectation step of the algorithm, each MNase-seq fragment is probabilistically assigned to nucleosome components according to the current locations, strengths, and subtype identities of the components. **D)** In the Maximization step of the algorithm, the various nucleosome properties are updated based on the current fragment assignments. **E)** Detailed illustration of how nucleosome properties are updated during EM iterations.

SEM runs in two major phases: nucleosome subtype discovery and nucleosome finding. During the first phase, SEM fits a GMM onto the fragment size distribution of the entire MNase-seq dataset to determine the Gaussian distribution parameters of each nucleosome subtype. The number of clusters, corresponding to the number of nucleosome subtypes, is an essential parameter of the model. It can be specified by the user according to prior knowledge of the biological sample, or SEM can automatically determine the best fit value using a Dirichlet Process Mixture Model (DPMM).

In the second phase, SEM uses a hierarchical GMM to compute the likelihood that each specific nucleosome is responsible for generating each MNase-seq fragment in the dataset. A Generalized Expectation Maximization (GEM) framework is used to calculate the latent assignment of MNase-seq fragments to nucleosomes and to estimate the various properties associated with each nucleosome (see **Methods**). Briefly, the Expectation step in the algorithm probabilistically assigns MNase-seq fragments to the nucleosome components that are most likely to have generated them based on their current locations and properties. Then the Maximization step updates the properties of the nucleosome components based on the best fit to the current fragment assignments. For example, a nucleosome component’s dyad location, occupancy, and fuzziness properties are re-estimated based on the mean midpoint location, weights, and variance of currently assigned MNase-seq fragments. Subtype identity is updated based on the size distribution of assigned fragments. The algorithm iterates through Expectation and Maximization steps until convergence.

To better reflect the known biological properties of nucleosomes, several priors are integrated into the model. First, a sparse prior on nucleosome occupancy eliminates components that don’t have sufficient numbers of fragments assigned, thereby encouraging solutions where each nucleosome is supported by MNase-seq data. Another sparse prior on subtype probability encourages each component to be a member of individual subtypes, which improves the interpretability of nucleosome subtype assignments. The physical exclusion of adjacent nucleosomes is also taken into consideration to ensure that neighboring nucleosomes do not overlap in the model.

### SEM accurately predicts conventional nucleosome properties

We first evaluated the performance of SEM in predicting conventional nucleosome properties, including nucleosome dyad location, nucleosome occupancy, and fuzziness, on both simulated and real MNase-seq datasets. To generate the simulated dataset, we took reference nucleosome dyad locations from yeast Jiang *et al*. 2009^10^, which is a compiled collection of nucleosome dyad locations from six datasets. The occupancy and fuzziness of each nucleosome were then computed using a H4 MNase-ChIP-seq dataset against the reference nucleosome dyads (see **Methods**). Background “noise” reads following a Poisson distribution were added globally to the simulated dataset to test the robustness of each algorithm.

SEM’s performance in this simulated dataset was evaluated against two other nucleosome calling packages: DANPOS and PuFFIN ^17,20^. All three packages were executed using their default settings (see **Methods**). The numbers of nucleosomes predicted by each package was similar to the number in the reference set (i.e., ∼60,000 nucleosomes). SEM and PuFFIN outperform DANPOS in predicting simulated nucleosome dyad locations (**Fig. 2A**). PuFFIN achieves the best estimates of simulated nucleosome occupancy, as evaluated by Pearson Correlation Coefficient (**Fig. 2D**). While SEM and DANPOS have similar correlations with the simulated occupancy, DANPOS has a general tendency to overestimate nucleosome occupancy (**Fig. 2D, middle panel**). SEM outperforms PuFFIN and DANPOS in estimating the simulated fuzziness levels, again evaluated by Pearson Correlation Coefficient (**Fig. 2E**).

**Figure 2:**
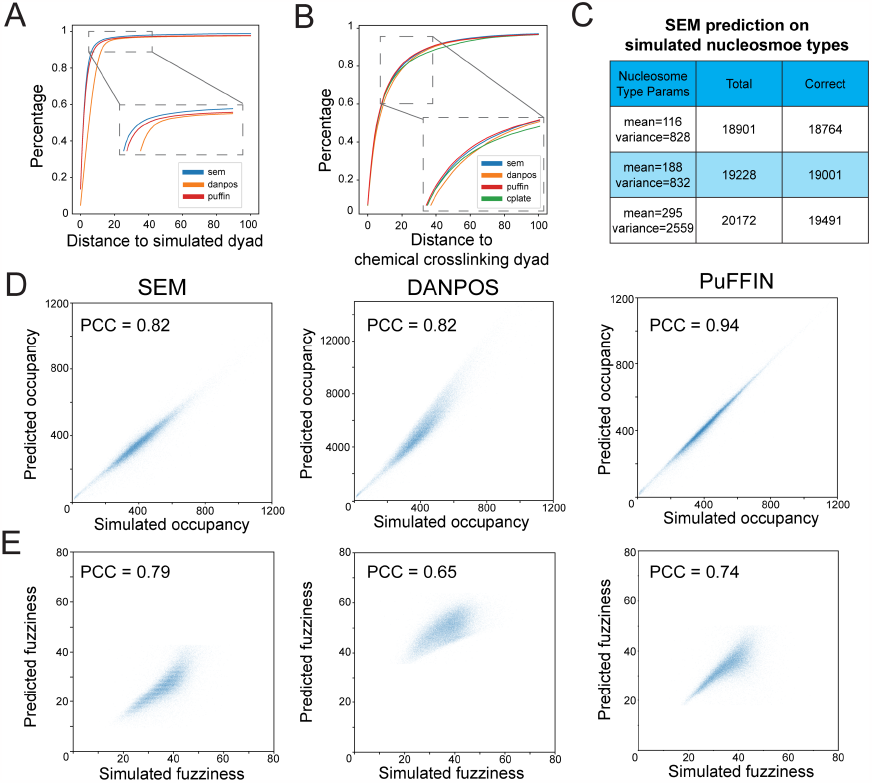
Comparison of SEM to existing nucleosome calling packages on common metrics. **A)** Cumulative percentage of distance between predicted nucleosome dyads and simulated nucleosome dyads in simulated MNase-seq data. SEM was executed twice with different occupancy thresholds (alpha=1 or alpha=5). **B)** Cumulative percentage of distance between predicted nucleosome dyad and chemical crosslinking dyad in real MNase-seq data (Voong et al., 2016). **C)** Number of total simulated nucleosomes and correctly predicted nucleosomes of each nucleosome subtype. **D)** Correlation between predicted occupancy and simulated occupancy. Each simulated nucleosome is assigned to the closest predicted nucleosome for the purposes of comparison. **E)** Correlation between predicted fuzziness and simulated fuzziness. Each simulated nucleosome is assigned to the closest predicted nucleosome for the purposes of comparison.

We further evaluated the performance of SEM using real MNase-seq data. Since no ground truth of nucleosome occupancy and fuzziness is available, our evaluation solely focused on predictions of nucleosome dyad locations. The nucleosome dyad locations obtained from the chemical crosslinking experiment were used as the “gold standard” due to its ability to specifically break DNA nucleotides near the dyad location. We included a fourth nucleosome calling package, Cplate, in this comparison, as predicted nucleosome dyad locations in the same dataset were available in the original Cplate publication ^18^. Notably, although SEM, DANPOS, and PuFFIN exhibited significant performance disparities in simulated datasets, all four software packages exhibited comparable performance when evaluated on the real MNase-seq dataset (**Fig. 2B**).

Additionally, we evaluated SEM’s ability to distinguish various nucleosome subtypes. A simulated MNase-seq dataset, where reads are sampled from three distinct nucleosome subtypes, was generated. SEM achieved high accuracy in correctly identifying the subtypes from which each simulated nucleosome was sampled (**Fig. 2C**).

To summarize, our results indicate that SEM performs comparably with existing nucleosome calling packages in predicting nucleosome dyad location. Although PuFFIN displays greater accuracy in predicting nucleosome occupancy, SEM outperforms it in predicting nucleosome fuzziness. Overall, SEM yields comparable performance to other nucleosome calling packages when it comes to conventional nucleosome metrics. In addition, SEM can precisely predict nucleosome subtypes in simulated data.

### Genome-wide detection of nucleosome subtypes in mouse embryonic stem cells

To assess whether SEM can distinguish different types of nucleosomes from MNase-seq data, we applied it to a low-dose MNase H2B ChIP-seq dataset from mESCs ^14^. The original study focused on characterizing the locations of so-called “fragile”, or MNase-sensitive, nucleosomes, which protect shorter DNA fragments than canonical nucleosomes under low dose MNase digestion ^14,21^. While the study found evidence for MNase-sensitive nucleosomes at TSSs and CTCF binding sites, the overlapping fragment size distributions of canonical nucleosomes and fragile nucleosomes makes it difficult to distinguish between these nucleosome subtypes at individual sites. The original study applied hard fragment length cut-offs (50-100bp, 101-140bp, 140-190bp) to discriminate different nucleosome subtypes, but these rigid thresholds give the appearance that distinct nucleosome subtypes overlap within the same regions (**Fig. 3C, S1A**). Our goal here is to assess whether SEM can more clearly discriminate nucleosome subtypes and add additional insights into the nature of the MNase-sensitive nucleosomes characterized in the original study.

**Figure 3:**
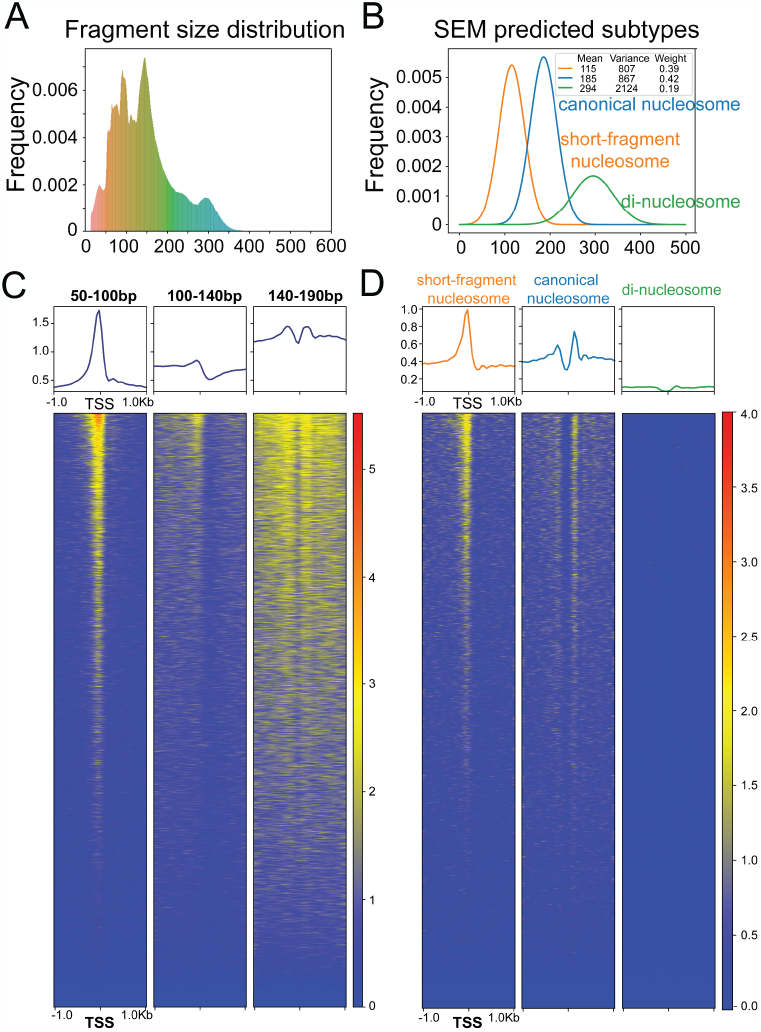
Subcategorization of SEM nucleosome subtypes reveals a population of accessible short-fragment nucleosomes. **A)** Fragment size distribution of the low dose MNase-ChIP-seq dataset. **B)** Fragment size distribution of each nucleosome subtype as determined by SEM. **C)** Heatmap and profile plot of MNase-seq fragments split by fragment size (50-100bp, 100-140bp, 140-190bp) around TSSs. **D)** Heatmap and profile plot of each SEM-defined nucleosome subtype around TSSs.

SEM detected three nucleosome subtypes in the low-dose MNase H2B ChIP-seq dataset. Based on their mean fragment sizes, we refer here to each subtype as: short-fragment nucleosomes (∼115bp mean size); canonical nucleosomes (∼185bp mean size); and di-nucleosomes (∼295bp mean size) (**Fig. 3A, B**). SEM discovered a total of 6,200,234 nucleosomes along the mouse genome. More than half exhibit mixed nucleosome subtype identity, indicating the highly dynamic nature of nucleosomes across the whole genome (**Fig. S1C, D**). In comparison with hard fragment length cut-offs, SEM’s detection of nucleosome subtypes more clearly separates nucleosomes belonging to each subtype at TSSs and CTCF sites (**Fig. 3D, S1B**). Specifically, short-fragment nucleosomes are enriched at the centers of TSSs and CTCF sites, while canonical nucleosomes are well-positioned in the flanking regions (**Fig. 3D, S1B**).

The enrichment of SEM-defined short-fragment nucleosomes at TSSs and CTCF sites mirrors that of the MNase-sensitive nucleosomes defined in the original study. However, only a small proportion of all SEM-defined short-fragment nucleosomes overlap TSSs or CTCF sites (**Fig. 4A**). We therefore asked whether SEM’s short-fragment nucleosome category consists of a homogenous class of MNase-sensitive nucleosomes or whether it encompasses multiple short-fragment nucleosome subtypes. To more clearly delineate the properties of each nucleosome subtype, we focus only on SEM-defined nucleosomes with occupancy greater than 5 reads and unambiguous subtype assignments (i.e., one of the nucleosome subtype probabilities >0.9).

**Figure 4:**
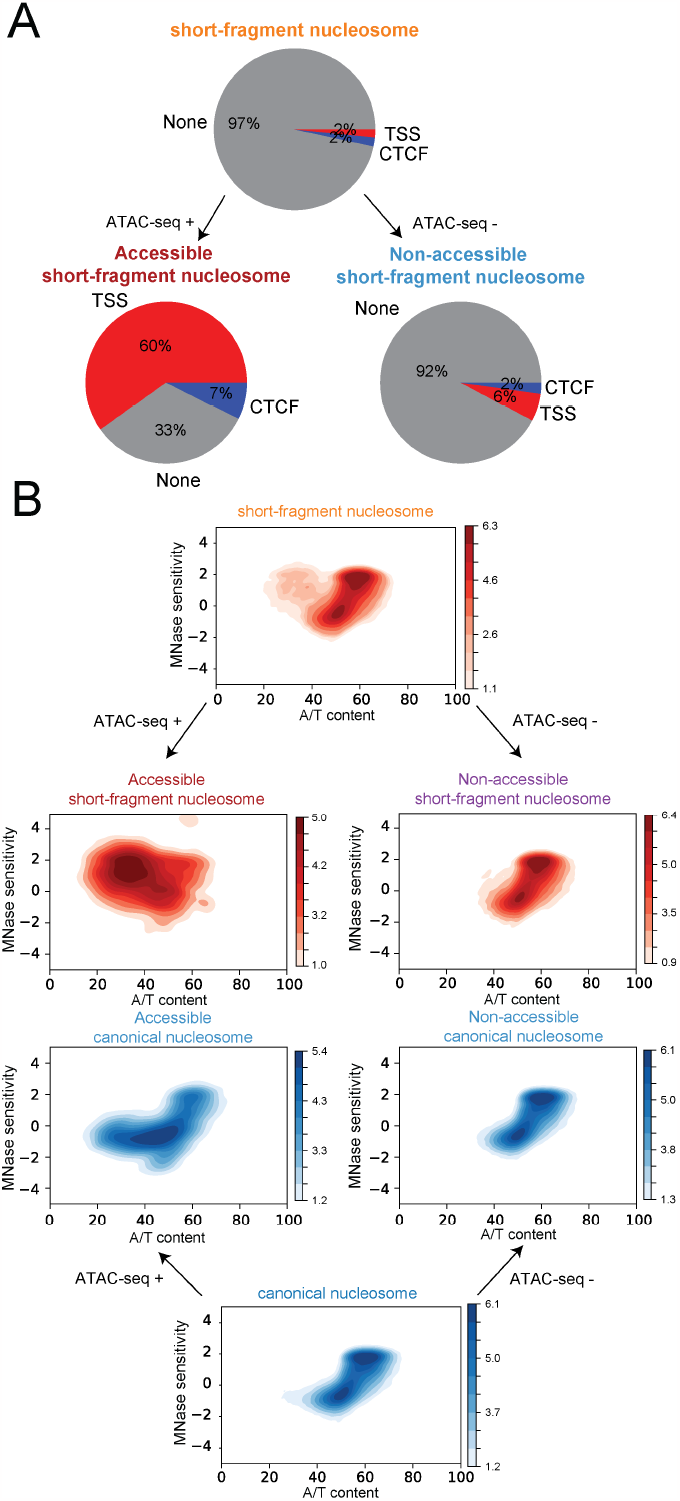
**A)** Percentages of short fragment nucleosome subtype and subcategories located at TSS regions (+/-500bp) and CTCF binding site. **B)** 2D density plots show the relationships between A/T content and MNase sensitivity of each nucleosome subtype and subcategory, color bar indicates the log density of nucleosomes. Nucleosome subcategories were obtained by examining if the nucleosome is fully overlapped by ATAC-seq peak (ATAC-seq +) or not (ATAC-seq -).

We first subcategorized SEM’s nucleosome subtypes according to their association with potential regulatory regions by overlapping with mESC ATAC-seq peaks ^22^. Two thirds of the SEM-defined accessible short-fragment nucleosomes overlap with TSSs or CTCF binding sites, while the non-accessible short-fragment nucleosomes are primarily located in distal regions (**Fig. 4A**). We then calculated the MNase sensitivity of each nucleosome subcategory by comparing the number of MNase-seq fragments from the low-dose MNase H2B ChIP-seq dataset with the number of fragments from a high-dose MNase-seq dataset ^14^ in the +/-50bp region surrounding each nucleosome dyad (**Fig. S2A**). Accessible short-fragment nucleosomes display high MNase sensitivity, consistent with the MNase-sensitive short fragment nucleosomes found in the original study. However, the non-accessible short-fragment nucleosomes do not display significantly higher MNase sensitivity than non-accessible canonical nucleosomes (**Fig. S2A**).

Since the non-accessible short-fragment nucleosomes do not display higher MNase sensitivity, we asked whether their shorter fragment lengths could be accounted for by MNase digestion biases towards A/T-rich sequences ^23^. We therefore compared the MNase sensitivity index for each nucleosome to the A/T content in the +/-50bp region surrounding each nucleosome dyad (**Fig. 4B**). While higher A/T content appears to correlate with higher MNase sensitivity at non-accessible nucleosome categories, the A/T content properties of non-accessible short-fragment nucleosomes are again similar to those of non-accessible canonical nucleosomes. However, non-accessible short-fragment nucleosomes display significantly higher A/T content at entry and exit sites compared with canonical nucleosomes (**Fig. S2B**). This suggests that the MNase bias towards digesting A/T-rich sequences at nucleosome flanks could be responsible for the shorter fragment distributions at this subset of short-fragment nucleosomes. In other words, there may not be any substantial difference between the non-accessible short-fragment nucleosomes and canonical nucleosomes as defined by SEM, other than a greater susceptibility to MNase biases at entry and exit sites.

In contrast to the non-accessible short fragment nucleosomes, the accessible short fragment nucleosomes display high MNase sensitivity despite being generally G/C rich (**Fig. 4B**). We confirmed the levels of MNase sensitivity using the MNase accessibility (MACC) score ^24^. The MACC score is the slope of the fitted line between fragment counts and logarithmic scaled MNase concentration in titrated MNase-seq experiments, where a high MACC score represents a highly MNase sensitive region. The MACC score is substantially higher at accessible short-fragment nucleosomes compared with random nucleosome positions (**Fig. S3A**). This indicates that SEM-defined accessible short-fragment nucleosomes represent a subpopulation of G/C-rich, MNase-sensitive short-fragment nucleosomes that is distinctly separated from the broader pool of short-fragment nucleosomes.

While the analyzed MNase-ChIP-seq experiment incorporated a H2B ChIP pulldown, it is possible that our detected short-fragment nucleosome subcategories result from MNase protection by non-nucleosomal regulatory proteins and artefactual crosslinking with neighboring nucleosomes. To address this potential concern, we examined nucleosome centering positioning (NCP) scores derived from a H4S47C-mediated chemical crosslinking approach ^16^. The accessible short-fragment nucleosomes display a clear peak in NCP scores, and nucleosome array phasing similar to canonical nucleosomes, providing orthogonal evidence for the presence of nucleosomes at the short-fragment nucleosome loci (**Fig. S3B**).

In summary, SEM identified three distinct nucleosome subtypes in mESCs from the low-dose MNase H2B ChIP-seq dataset. By integrating chromatin accessibility data, the SEM-defined short-fragment nucleosomes were further subdivided into two subcategories. Accessible short-fragment nucleosomes display several properties corresponding to the previously proposed fragile nucleosomes, including high MNase sensitivity and enrichment at TSSs and CTCF binding sites.

### Accessible short-fragment nucleosomes associate with distinct regulatory activities

Most previous studies of fragile or MNase-sensitive nucleosomes have focused on their occurrence at TSS and CTCF binding sites ^14,16^. As shown above, SEM-defined accessible short-fragment nucleosomes are highly enriched at these regions (**Fig. 4A**). However, a substantial portion (33%) of SEM’s accessible short fragment nucleosome do not overlap either TSSs or CTCF binding sites (**Fig. 4A**). To investigate the types of locations occupied by non-TSS/CTCF accessible short fragment nucleosomes, we overlapped these nucleosomes with candidate cis-regulatory elements (cCREs) from ENCODE SCREEN project ^25^ (**Fig. 5A**). Notably, 17% of the accessible short fragment nucleosomes overlap with either proximal or distal enhancer-like signatures (pELSs, dELSs). These results suggest accessible short-fragment nucleosomes are broadly associated with cis-regulatory elements.

**Figure 5:**
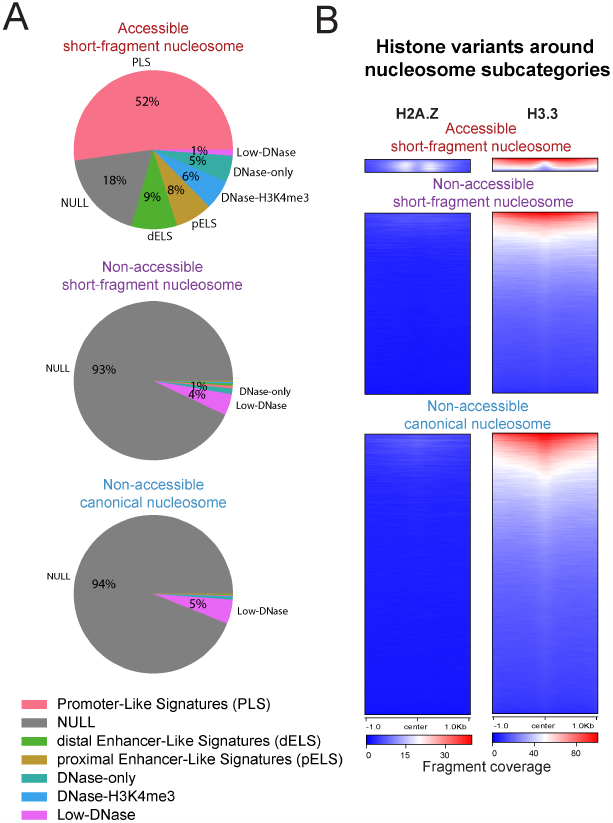
**A)** Pie chart summarizing the overlap between SEM-defined nucleosome subcategories and ENCODE SCREEN cCRE types. **B)** Heatmap displaying the enrichment of histone variants H2A.Z and H3.3 around nucleosome dyad locations.

The association between accessible short-fragment nucleosomes and cis-regulatory elements suggests a possible role in transcriptional regulation. Therefore, we investigated whether accessible short-fragment nucleosomes overlap the occupancy of factors involved in nucleosome composition. We also assessed the overlap with histone modifications associated with regulatory activities in mES cells. A previous report in Drosophila suggested that a set of H3.3/H2A.Z containing nucleosomes is enriched at regulatory regions and is unstable under high salt conditions ^26^. Thus, we examined the enrichment of both histone variants around nucleosome subcategories. Indeed, H3.3, but not H2A.Z, is enriched at accessible short-fragment nucleosome loci (**Fig. 5B**). Active histone modifications, including H3K27ac, H3K4me3, H3K9ac, and H4ac, were enriched at sites adjacent to accessible short-fragment nucleosomes (**Fig. S4**), while repressive marks, including H3K27me3 and H2AK119ub, do not show enrichment at accessible short-fragment nucleosome sites. Finally, several chromatin remodelers are enriched at accessible short-fragment nucleosome dyad locations, including Smarca4, Chd4, Chd6, Chd8, and Ep400 (**Fig. S5**). These chromatin remodelers are known to influence the contact between DNA and the nucleosome ^27^. Ep400 has also been reported to deposit H3.3 at promoter and enhancer regions ^28^, which may correspond to the H3.3 enrichment observed at accessible short fragment nucleosome sites. Therefore, the enrichment of chromatin remodelers at accessible short fragment nucleosomes suggests a mechanism for their destabilization.

In summary, accessible short-fragment nucleosomes defined by SEM are not restricted to TSSs and CTCF binding sites and correspond to sites of specific chromatin remodeler enrichment. While the regulatory roles played by these atypical nucleosomes require further investigation, our analyses demonstrate the efficacy of SEM in characterizing diverse nucleosome types.

## DISCUSSION

In recent years, there has been growing interest in nucleosome subtypes and compositional variants beyond canonical nucleosomes ^12,13,15,29^. Here, we introduced SEM, a new nucleosome calling framework capable of distinguishing various nucleosome subtypes from a single MNase-digested sequencing experiment. When compared to other nucleosome calling packages, SEM exhibits comparable performance on conventional nucleosome calling metrics while uniquely providing an automatic annotation of nucleosome subtype identity.

Using SEM on a low-dose MNase H2B ChIP-seq dataset from mESCs, we identified a nucleosome subtype that protects shorter DNA fragments than canonical nucleosomes. A further integration of ATAC-seq data classified the short-fragment nucleosomes into two subcategories, which we labeled accessible short-fragment nucleosomes and non-accessible short-fragment nucleosomes. Although SEM-defined non-accessible short fragment nucleosomes are likely the result of MNase digestion bias at entry and exit sites, it is worth noting that this nucleosome subcategory could be overlooked in traditional MNase-seq experiments conducting mono-nucleosome size selection. Thus, MNase-seq experiments that omit mono-nucleosome size selection, in combination with SEM analysis, could more accurately characterize nucleosome positions across the genome.

SEM-defined accessible short-fragment nucleosomes share a similar distribution to previously reported fragile nucleosomes at TSS and CTCF sites. However, SEM’s ability to identify short-fragment nucleosomes genome-wide shows that accessible short-fragment nucleosomes are not restricted to TSS and CTCF sites but are also present in enhancer regions. Previous studies have suggested that histone variants H3.3 and H2A.Z could regulate nucleosome stability or lead to nucleosome fragility ^13,26,30^. We find an enrichment of H3.3, but not H2A.Z, at accessible short-fragment nucleosome sites. Accessible short-fragment nucleosomes are also associated with active histone marks but not repressive marks. Accessible short-fragment nucleosomes in mESCs are co-enriched with the chromatin remodelers Smarca4, Chd4, Chd6, Chd8, and Ep400, suggesting that this nucleosome subcategory represent hotspots of chromatin remodeler activity. Further work remains to investigate whether SEM-defined accessible short-fragment nucleosomes play distinct regulatory roles in mESCs. However, our analyses demonstrate that SEM’s genome-wide probabilistic approach to nucleosome subtype calling provides clear advantages over the typically employed fragment length threshold approaches.

## METHODS

The SEM software package is available on GitHub (https://github.com/YenLab/SEM) and Bioconda (https://anaconda.org/bioconda/sem).

### Nucleosome subtype characterization

To model different nucleosome subtypes present in a given input dataset, we first apply a Gaussian Mixture Model (GMM) to the overall fragment size distribution. This step can be processed with or without a user-defined number of subtypes. If the number of types has been defined, a finite GMM is used on the fragment size distribution to find the parameters of each type. Otherwise, a Dirichlet Process Mixture Model (DPMM) is employed, and SEM will automatically determine the number of subtypes.

### SEM probabilistic model

SEM is based on a hierarchical Gaussian Mixture Model that describes the likelihood of observing a set of MNase-seq fragments from a set of nucleosomes. Each nucleosome contributes a distribution of reads surrounding its genomic position to the overall mixture of reads. We assume that fragment locations are independently conditioned on the dyad location, fuzziness, and subtype mixture probabilities of their underlying nucleosomes.

SEM performs nucleosome discovery by finding the set of nucleosomes that maximizes the penalized likelihood of the observed MNase-seq fragments. First, we assume there are *K* nucleosome fragment size subtypes, where each subtype is a Gaussian distribution with mean *Ψ*= *ψ*_1_, …, *ψ*_*K*_, variance *Φ* = *ϕ*_1_, …, *ϕ*_*K*_, and weight *Ω* = *ω*_1_, …, *ω*_*K*_. Thus, the expected size distribution of fragments from each nucleosome follows a mixture of the subtype distributions, conditioned on their subtype probabilities. Throughout the whole genome, we consider *N* MNase-seq fragments that have been mapped to genomic locations *E* = *ϵ*_1_, …, *ϵ*_*N*_ with size *H* = *η*_1_, …, *η*_*N*_, and *M* potential nucleosomes at genomic locations *U* = *μ*_1_, …, *μ*_*M*_ with fuzziness *Θ* = *θ*_1_, …, *θ*_*M*_ and subtype mixture probabilities *T* = *τ*_1_, …, *τ*_*M*_ = [*τ*_1,1_, …, *τ*_1,*K*_,], …,[*τ*_*M*,1_, …, *τ*_*M,K*_,]. The locations of fragments from nucleosome *m* follow a Gaussian distribution, where the Gaussian mean equals the nucleosome dyad location *μ*_*m*_ and the variance equals the fuzziness parameter *θ*_*M*_. We represent the latent assignment of fragments to nucleosomes that caused them as *Z* = *z*_1_, …, *z*_*N*_, where *z*_*i*_ = *j* where *j* is the index of the nucleosome whose dyad is at position *μ*_*j*_ that generates fragment *i*.

The conditional probability of fragment *r*_*i*_ being generated from nucleosome *j* located in *μ*_*j*_ with mixture probability ρ_*j*_ and fuzziness *θ*_*j*_ is:

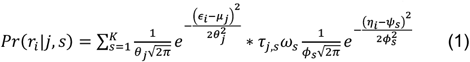

where *τ*_*j,s*_ represents the probability of nucleosome *j* belonging to fragment size type *s*.

The probability of a fragment is a convex combination of possible nucleosomes:

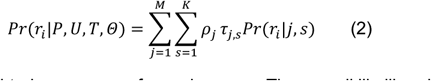

where *ρ* represents the weighted occupancy of a nucleosome. The overall likelihood of the observed set of fragments is then:

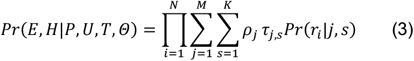

We incorporate several biological assumptions in the form of priors on the parameters. We place a sparseness-promoting negative Dirichlet prior *α* on the nucleosome weighted occupancy *π*, based on the assumption that each nucleosome should have sufficient numbers of assigned fragments to support its existence in the model:

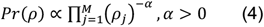

We also incorporate an additional prior to encourage nucleosomes to choose less ambiguous subtype identities. To do so, we place a Dirichlet prior *β* on the nucleosome type mixture probability *τ*.

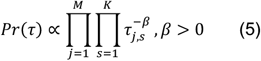

In summary, the complete-data log posterior is as follows:

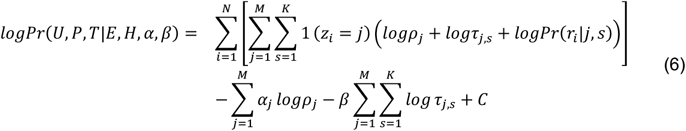

### Expectation Maximization (EM)

We initialize mixing probabilities *ρ* with uniform probabilities, *ρ*_*j*_= 1/*M, j* = 1, …, *M*. At the E step, we calculate the relative responsibility of each nucleosome subtype per nucleosome in generating each fragment as follows:

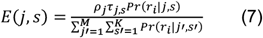

The maximum posterior probability (MAP) estimation of *ρ, τ* is as follows:

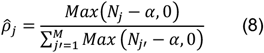

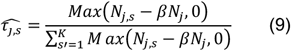

where *N*_*j*_ represents the number of fragments assigned to nucleosome *j, N*_*j,s*_ represents the number of fragments assigned to nucleosome *j* of type *s*. The value of *α* is the minimum number of MNase-seq fragments required to support a nucleosome surviving in this round. In the first five rounds of EM, sparse priors are set to their minimum value (1 for *α* and 0 for *β*) to avoid elimination of true nucleosome due to bad initialization. Then sparse priors are gradually increased in the following five rounds until reaching the default or user defined values. The default values of *α* and *β* are 1 and 0.05, respectively.

MAP values of *μ*_*j,s*_ are determined by finding the location with the maximized probability in +/-50bp flanking sites of the nucleosome dyad. If the maximization step results in two components sharing the same weighted positions, they are combined in the next iteration of the algorithm.

Considering the efficiency of the algorithm, the fuzziness and the subtype mixture probabilities of each nucleosome are updated every two rounds. In the first several rounds, prior *α* and *β* is multiplied by an annealing factor according to the number of finished rounds to avoid too early elimination of nucleosomes. As EM proceeds, the log likelihood increases after each iteration, and convergence is defined when log likelihood increases less than 1% compared to previous iteration.

### Exclusion zone

Each nucleosome protects DNA and should sterically exclude other nucleosomes at the same position. For example, canonical nucleosomes protect 147bp DNA, which means the minimum distance between two canonical nucleosomes should be larger than 147bp. Thus, when there is no nucleosome eliminated during an iteration after sparse prior fully incorporated, an extra step will be taken to remove nucleosomes which are too close to adjacent nucleosomes. Each time the overlapped nucleosome with the lowest responsibility (i.e., occupancy) will be removed, until all nucleosomes have a large enough spacing to each other. The exclusion zone is currently set to 127bp to avoid false removal of nucleosomes due to the inaccurate prediction of nucleosome dyad.

### MNase-seq data simulation

To simulate the MNase-seq dataset, we first took the nucleosome dyad location maps from ^10^ as a reference. MNase H4 ChIP-seq datasets from SRR3649286, SRR3649291 ^21^ were used to infer the nucleosome occupancy and fuzziness for simulation. Specifically, we used the MNase-seq fragments within +/-73bp around each nucleosome dyad to compute the occupancy and fuzziness. Simulated MNase-seq fragments were then generated given the computed nucleosome metrics, distributed following a Gaussian with mean (nucleosome dyad), variance (nucleosome fuzziness), and weight (nucleosome occupancy) parameters set from the above data. Background noise fragments following a Poisson Distribution were added to the simulated dataset to test the robustness of each algorithm. We used background noise ratio 0.05 in this study. The Poisson mean *λ* is determined by the ratio of background noise among the whole dataset using the formula:

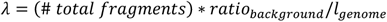

### Nucleosome calling on MNase-seq datasets

On both simulated and real datasets, SEM, DANPOS, and PuFFIN were run under default setting. Cplate predictions on real MNase-seq dataset were taken from reference ^18^.

### Determining nucleosome subcategories in mESCs

SEM was run on low-dose MNase H2B ChIP-seq dataset (SRR2034510, SRR2034511) from ^14^ under default settings with the number of nucleosome subtypes set as 3. Nucleosomes are then filtered such that occupancy is greater than 5 reads and one of the nucleosome subtype probabilities > 0.9. The nucleosome subcategories were determined by overlapping the filtered nucleosomes with ATAC-seq peaks. To ensure the whole nucleosome is located within an accessible region, we required the dyad location of accessible nucleosome has to be at least 100bp away from the boundary of the ATAC-seq peak. The remaining nucleosomes are deemed non-accessible.

### Data processing

All fastq files were first trimmed by trimmomatic ^31^ with the options “LEADING:3 TRAILING:3 SLIDINGWINDOW:4:15 MINLEN:36 {adapter_file}:2:30:10*K* using the corresponding adapters, then trimmed fastq files were mapped to the genome by bowtie2 ^32^. Mapped bam files were filtered by the criteria that MAPQ of both reads in a pair>=10. ATAC-seq peak calling was performed by using Genrich (https://github.com/jsh58/Genrich) with ATAC-seq mode enabled by “-j” option.

## ACKNOWLEDGEMENTS

This work was supported by NIH R35-GM144135 and National Science Foundation DBI CAREER 2045500 (to S.M.). Any opinions, findings and conclusions or recommendations expressed in this material are those of the authors and do not necessarily reflect the views of the National Science Foundation. This work was initiated by J.Y. as a Master’s student in K.Y.’s lab at the Department of Developmental Biology, School of Basic Medical Sciences, Southern Medical University, Guangzhou, China. Work by the Yen lab is supported by the National Natural Science Foundation of China (grants 31522031 & 31571526). The authors thank the members of the Center for Eukaryotic Gene Regulation at Penn State for helpful feedback and discussions.

## Notes

### Competing Interest Statement

The authors have declared no competing interest.

