## Supplemental Figures for "SEM: sized-based expectation maximization for characterizing nucleosome positions and subtypes"

### SUPPLEMENTARY FIGURES

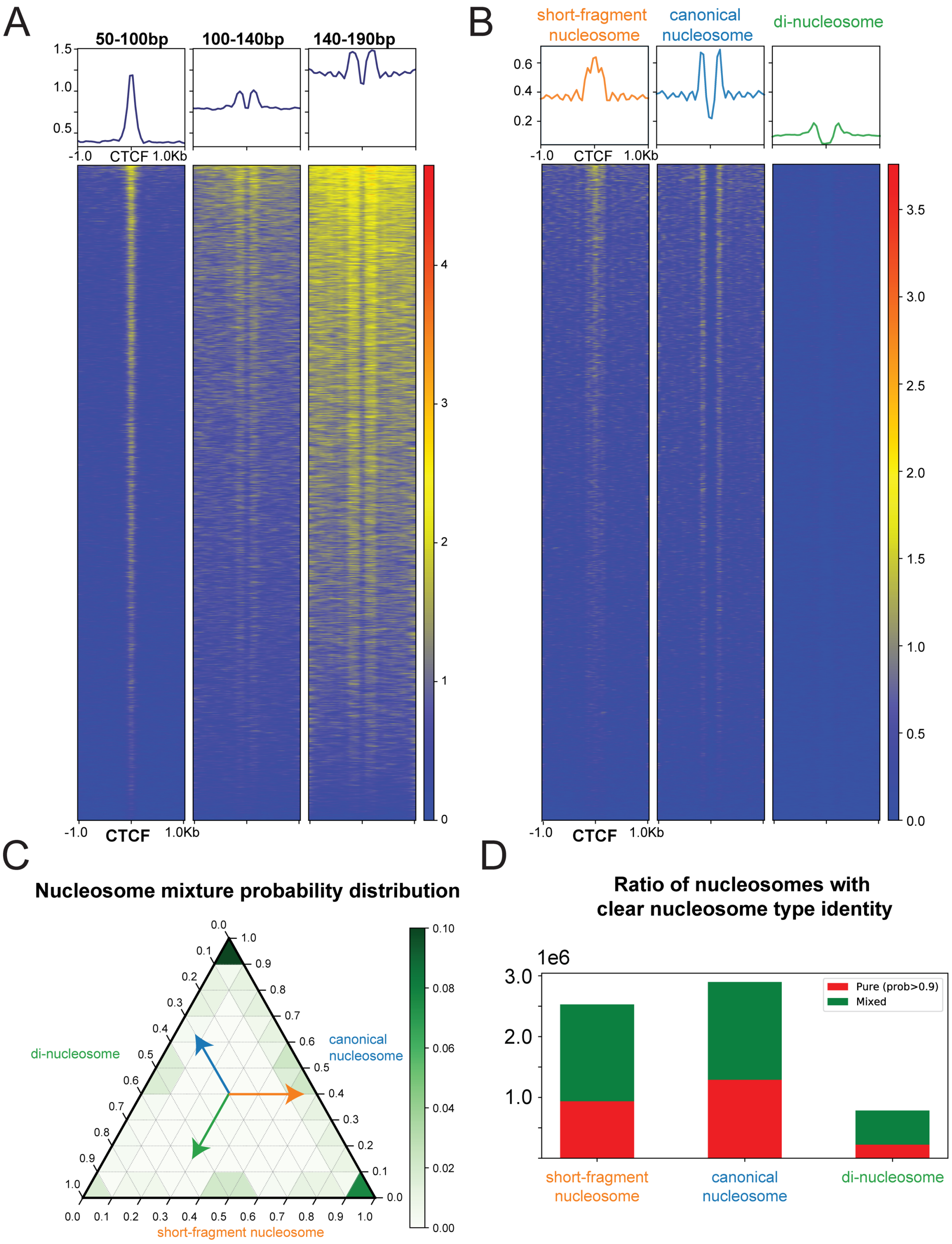


**Figure S1: A)** Heatmap and profile plot of MNase-seq fragments split by fragment size (50-100bp, 100-140bp, 140-190bp) around CTCF binding sites**. B)** Heatmap and profile plot of each nucleosome subcategory around CTCF binding sites. **C)** Ternery plot showing the distribution of nucleosome mixture type probabilities. **D)** Bar plot shows the proportion of mixed and "pure" (subtype probability > 0.9) nucleosomes for each of the nucleosome subtypes.

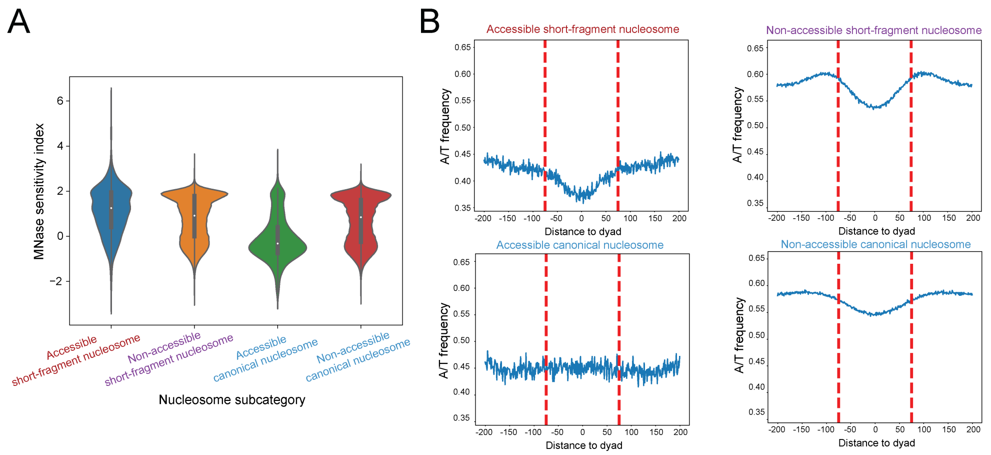


**Figure S2: A)** Violin plot shows the distribution of MNase sensitivity index in each nucleosome subcategory. **B)** A/T nucleotide frequency around each nucleosome subcategory, red dash lines mark the nucleosome entry/exit sites (+/-75bp).

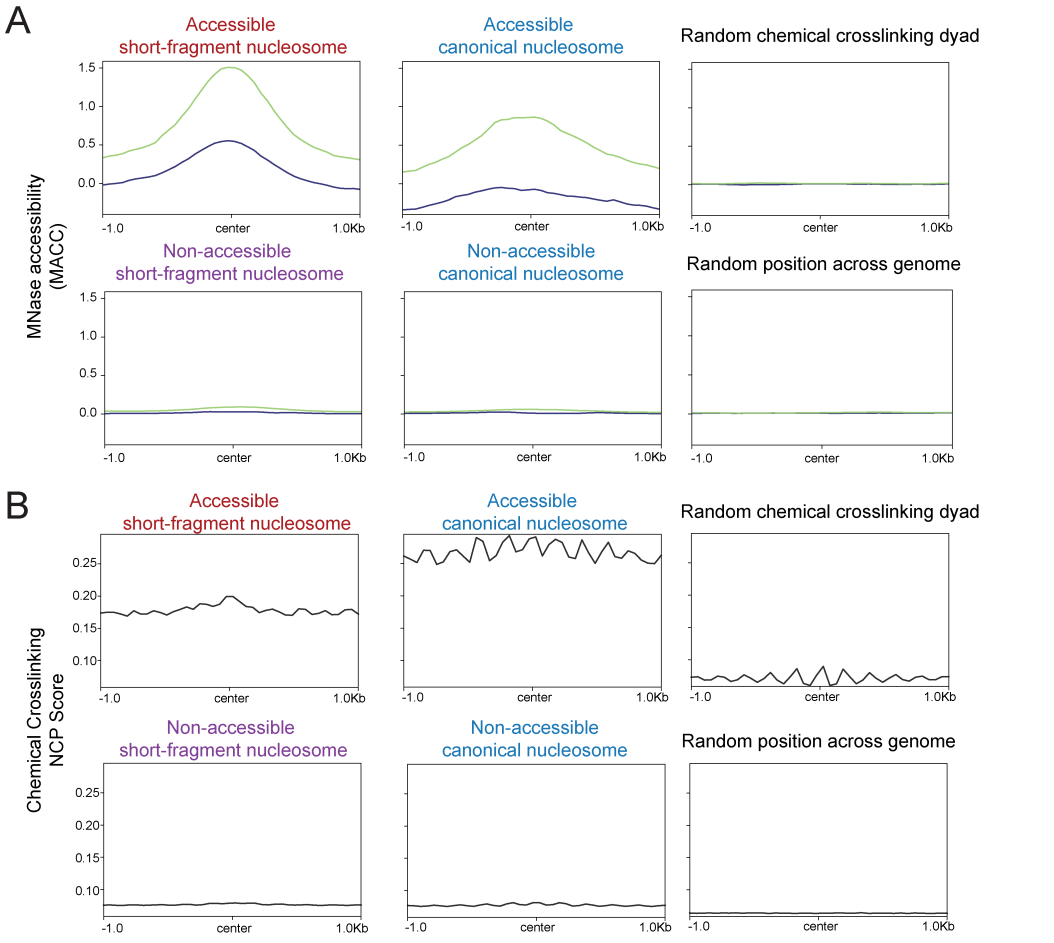


**Figure S3: A)** Profile plot of MACC value around nucleosome dyad locations for each nucleosome subcategory. **B)** Profile plot of Chemical Crosslinking NCP score around nucleosome dyad locations for each nucleosome subcategory.

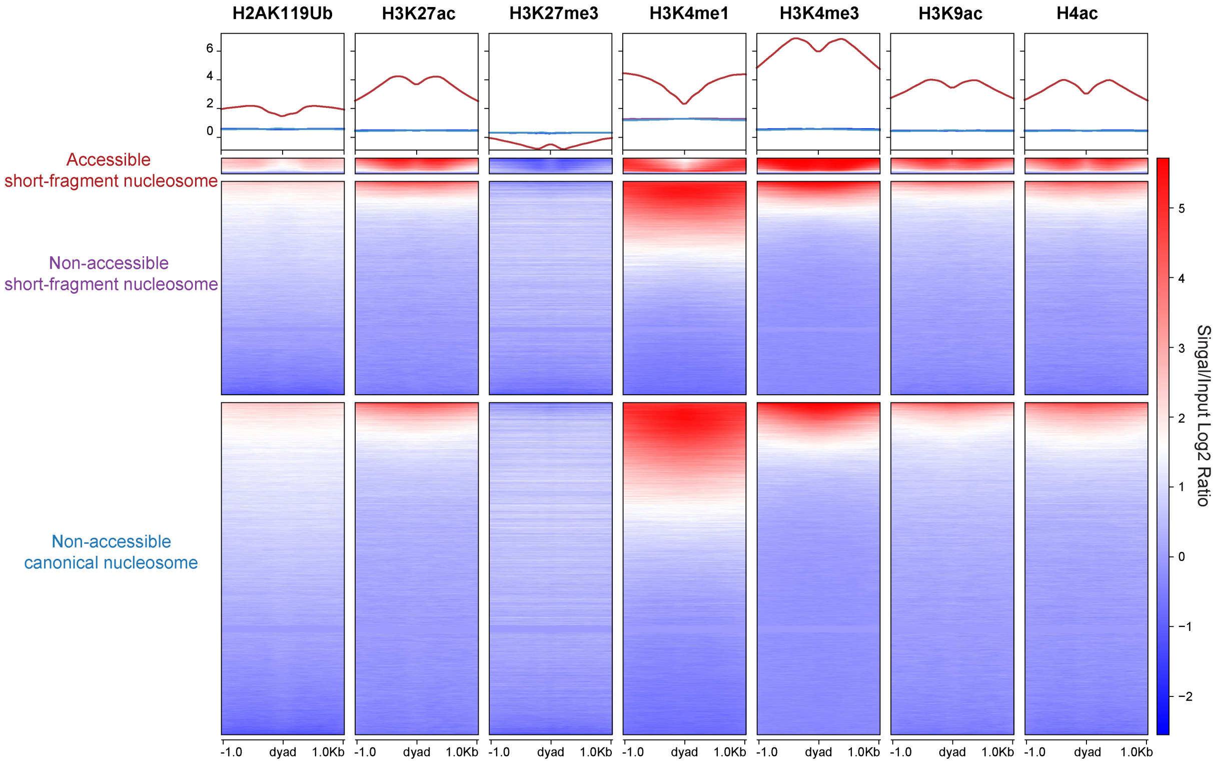


**Figure S4:** Heatmap and profile plot of histone modifications around nucleosome dyad locations.

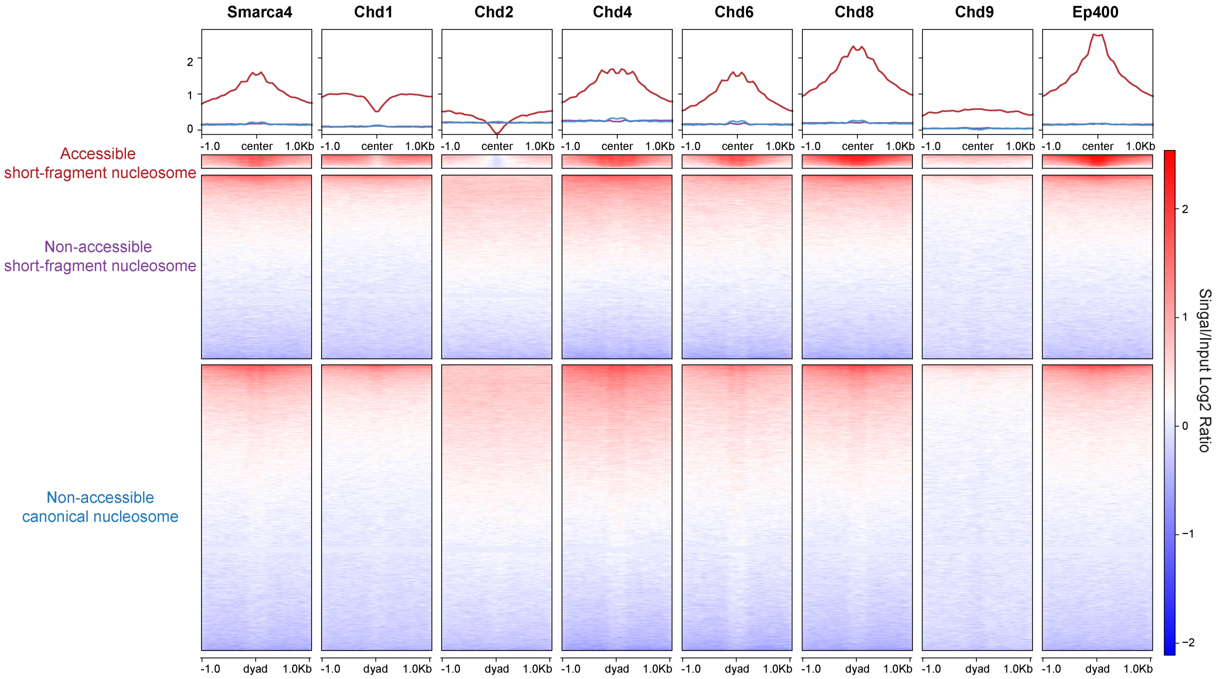


**Figure S5:** Heatmap and profile plot of chromatin remodelers around nucleosome dyad locations.
